# Affective Symptoms in Pregnancy are Associated with the Vaginal Microbiome

**DOI:** 10.1101/2024.04.12.589254

**Authors:** Kristin Scheible, Robert Beblavy, Michael B. Sohn, Xing Qui, Ann L. Gill, Janiret Narvaez-Miranda, Jessica Brunner, Richard K. Miller, Emily S. Barrett, Tom G. O’Connor, Steven R. Gill

**Affiliations:** Department of Microbiology and Immunology, University of Rochester School of Medicine and Dentistry, Rochester, New York, USA; Department of Pediatrics, University of Rochester School of Medicine and Dentistry, Rochester, New York, USA; Department of Biostatistics and Computational Biology, University of Rochester School of Medicine and Dentistry, Rochester, New York, USA; Department of Obstetrics and Gynecology, University of Rochester School of Medicine and Dentistry, Rochester, New York, USA; Biostatistics and Epidemiology, Rutgers School of Public Health, Piscataway, New Jersey, USA; Environmental and Occupational Health Sciences Institute, Rutgers University, Piscataway, New Jersey, USA; Department of Psychiatry, University of Rochester School of Medicine and Dentistry, Rochester, New York, USA; Department of Neuroscience, University of Rochester School of Medicine and Dentistry, Rochester, New York, USA; Wynne Center for Family Research, University of Rochester School of Medicine and Dentistry, Rochester, New York, USA

**Keywords:** Affective symptoms, vaginal microbiome, community state type, cortisol, sex hormones, socio-demographic

## Abstract

Composition of the vaginal microbiome in pregnancy is associated with adverse maternal, obstetric, and child health outcomes. Identifying the sources of individual differences in the vaginal microbiome is therefore of considerable clinical and public health interest. The current study tested the hypothesis that vaginal microbiome composition during pregnancy is associated with an individual’s experience of affective symptoms and stress exposure. Data were based on a prospective longitudinal study of a diverse and medically healthy community sample of 275 mother-infant pairs. Affective symptoms and stress exposure and select measures of associated biomarkers (diurnal salivary cortisol, serum measures of sex hormones) were collected at each trimester; self-report, clinical, and medical records were used to collect detailed data on socio-demographic factors and health behavior, including diet and sleep. Vaginal microbiome samples were collected in the third trimester (34-40 weeks) and characterized by 16S rRNA sequencing. Identified taxa were clustered into three community state types (CST1-3) based on dissimilarity of vaginal microbiota composition. Results indicate that depressive symptoms during pregnancy were reliably associated with individual taxa and CST3 in the third trimester. Prediction of functional potential from 16S taxonomy revealed a differential abundance of metabolic pathways in CST1-3 and individual taxa, including biosynthetic pathways for the neuroactive metabolites, serotonin and dopamine. With the exception of bioavailable testosterone, no significant associations were found between symptoms- and stress-related biomarkers and CSTs. Our results provide further evidence of how prenatal psychological distress during pregnancy alters the maternal-fetal microbiome ecosystem that may be important for understanding maternal and child health outcomes.

**Importance:** Prenatal affective symptoms and stress are associated with maternal, obstetric, and child health outcomes, but the mechanisms underlying these links and their application to intervention remain unclear. The findings from this investigation extend prior microbiome-oriented research by demonstrating that the maternal vaginal microbiome composition has a biologically plausible mechanistic link with affective symptoms that also suggest additional clinical applications for assessment and intervention.

## INTRODUCTION

Affective symptoms and stress in pregnancy are reliably associated with perinatal, obstetric and child health outcomes^1-3^; the underlying mechanisms are under intense scrutiny because of the potential applications to clinical practice and public health. Prenatal (vaginal) microbiome composition is a plausible candidate, but there are few studies examining how the vaginal microbiome in pregnancy responds to exposures and health behaviors, such as affective symptoms and stress. This relationship is particularly understudied among women with normal pregnancy risk, for whom the results may have the strongest public and clinical health impact. The current study contributes novel and significant findings to this research; specifically, we test the hypothesis that maternal affective symptoms and stress during pregnancy are associated with vaginal microbiome composition in a prospective longitudinal study of a diverse, well-characterized, and medically healthy sample.

Most investigations into the role of the microbiome in affective symptoms and stress of pregnant and non-pregnant women have been focused on the gut microbiota^4,5^. However, with significant differences in microbiota composition, function, and physiological niche, neither the patterns of associations nor the putative biological pathways from these efforts directly extend to the vaginal microbiome. Research findings linking the vaginal microbiome, affective symptoms, and plausible biological pathways are rare. Some of the limited available evidence derives from pre-clinical studies in animal models, which suggest that experimentally induced stress decreases vaginal microbiome diversity ^6,7^. More recent investigations of psychological and other exposures on the vaginal microbiome in pregnancy-age and pregnant individuals have shown an association between psychosocial stress and the vaginal microbiota, including decreased abundance of beneficial bifidobacterial species and increased bacterial vaginosis communities^4,8-11^. Studies in humans that model joint effects of psychosocial stress and vaginal microbiota in high-risk pregnancies showed increased risk for preterm birth^10,12^.

Research on the link between affective symptoms and stress and the vaginal microbiome may also clarify if the vaginal microbiome may be a pathway through which prenatal distress shapes child health outcomes ^6,9,13-19^. Colonization and temporal development of the infant gut microbiota in early life is initiated by mother-to-infant transfer of maternal gut and vaginal microbiota at birth^20-22^. Disruption of these interactions due to perturbation of the colonizing vaginal microbiota associated with maternal affective symptoms have the potential for a sustained impact on early life neurodevelopment and occurrence of affective symptoms in children^20,23-26^. This ability for the maternal microbiome to directly or indirectly impact fetal and child neurodevelopment is supported by animal studies^6,16,27^. A primary goal of this current work is to establish plausible biological pathways in the maternal vaginal microbiome, with affective symptoms and their associated biomarkers as primary variables in a healthy, normative risk pregnant cohort^28,29^. In addition to assessing maternal prenatal microbiome in relation to clinical measures of affective symptoms and stress, we also consider the degree to which microbiome composition associates with several biomarkers of these clinical measures. These findings will guide our future efforts to establish biological pathways in the early infant early life microbiome-gut-brain axis that influence affective symptoms and stress^6,16,29-35^.

## METHODS

### Study Overview and Sample

The current analysis is based on data from a prospective longitudinal cohort study of prenatal influences on child health outcomes based in Rochester, NY; “Understanding Pregnancy Signals and Infant Development” (UPSIDE)^36^ which is part of the NIH Environmental influences on Child Health Outcomes program^37^. The study was approved by the University of Rochester Research Subjects Review Board; all participants provided written informed consent. Participants were compensated for each research visit and were provided transportation if needed. For the UPSIDE cohort, women in their first trimester of pregnancy were recruited from obstetric clinics affiliated with the University of Rochester between December 2015 and April 2019. Eligibility criteria were age 18 or older, singleton pregnancy, no known substance abuse problems or a history of psychotic illness, ability to communicate in English, not greater than normal medical risk. Women with significant medical morbidities and endocrine disorders (e.g., polycystic ovary syndrome) or obstetric problems were excluded. Of the 326 women who were enrolled in the first trimester, 18 were excluded for pregnancy loss or heightened pregnancy risk (e.g., multiple pregnancy, miscarriage, medical screen failure). Of the remaining 308 participants, 275 had sufficient sample volume collection for microbiome analysis.

Study visits, conducted in a private clinic room, consisted of extensive biospecimen and questionnaire data collection, which were supplemented with health information abstracted from the medical record. In each trimester, women completed questionnaires assessing affective symptoms; life events stress was collected at trimester 3 and referred to the past year. Depressive symptoms were assessed via the Edinburgh Postnatal Depression Scale (EPDS), a well-validated and widely-used 10-item scale^38^ that is also validated for use in prenatal populations. A cut-off of ≥13^39^ has been used in previous literature to indicate possible clinical depression. Anxiety symptoms were self-reported using the Penn State Worry Questionnaire (PSWQ). In the third trimester, Stressful Life Events (SLE) were reported using an adapted version of the Inventory of Ranked Life Events for Primiparous and Multiparous Women; this 26-item scale measures stressful life events that occurred in the past year^40^. SLE was tallied as total number of events endorsed. Due to the right-skewed distribution of the total number of events, the scale was re-scored using a cutoff at 5, i.e., values ranged from 0 to 5 or more. Clinical covariates considered in the model included pre-pregnancy body mass index (BMI), antibiotic exposure within four weeks prior to vaginal swab collection, maternal self-identified race, parity, maternal age and fetal sex^36,41,42^.

### Diurnal cortisol assessment

Diurnal cortisol was self-collected by study participants following a standard passive drool protocol^43^. Samples were collected at home at five points across a single day at each trimester (at wake, 45 minutes post-wake, 2.5 hours, 8 hours, and 12 hours post-wake), for a total of 15 samples across gestation. Samples were stored in their collection tubes at -80°C until analysis. Cortisol was assayed using kits from Salimetrics, LLC (Carlsbad, CA, cat# 1-3002) following manufacturer instructions. Average intra- and inter-assay coefficients of variation were 2.40% and 11.75%. Area Under the Curve (AUC) with reference to ground is often utilized to reflect total daily cortisol load, with trapezoidal approximations estimating the total cortisol output of the system^44^. AUC calculations were limited to participants who had 4 or 5 viable daily samples including the early morning/wake-up sample.

### Sex steroid hormone measurement

Maternal and cord blood serum sex steroid hormone concentrations were measured at the Endocrine and Metabolic Research Laboratory at the Lundquist Institute at Harbor-UCLA Medical Center. Estrone (E1), E2, estriol (E3), total testosterone (TT) and free testosterone (fT) were measured in maternal serum. In cord blood we measured TT, fT, androstenodione (A4), and dehydroepiandrosterone (DHEA)^45,46^. Hormones were measured by LC-MS/MS using a Shimadzu HPLC system (Columbia, MD) interfaced with an Applied Biosystems API5500 LC-MS/MS (Foster City, CA). TT was measured with Turbo-Ion-Spray source in the positive ionization mode. Measurement of fT was done with equilibrium dialysis using labeled testosterone^45^. Concentrations below the LOD and missing values for E3 in the first trimester (n=25) were imputed with the LOD/√2.

### Vaginal sample collection

To obtain the vaginal sample, a sterile Catch-All™ sample collection swab was placed at the vaginal introitus immediately posterior to the hymenal ring. The swab was rotated in a circular motion five times along the lumen, removed from the vagina and immediately placed in 750 μL of sterile phosphate buffered saline, stored on ice for no more than 2 hrs. and transferred to -80°C for storage prior to DNA extraction.

### Genomic DNA extraction, PCR amplification and sequencing

Total genomic DNA was extracted from the vaginal samples using a modification of the ZymoResearch Quick-DNA Fecal/Soil Microbe Miniprep Kit (Zymo Research, Irvine, CA) and FastPrep mechanical lysis (MPBio, Solon, OH). 16S ribosomal RNA (rRNA) was amplified with Phusion High-Fidelity polymerase (New England Biolabs, Ipswich, MA) and dual indexed primers (319F: 5′ ACTCCTACGGGAGGCAGCAG 3′; 806R: 3′ ACTCCTACGGGAGGCAGCAG 5′) specific to the V3-V4 hypervariable regions^47^. Amplicons were pooled and paired-end sequenced on an Illumina MiSeq (Illumina, San Diego, CA) in the University of Rochester Genomics Research Center. Each sequencing run included a mock community of *Staphylococcus aureus, Lactococcus lactis, Porphyromonas gingivalis, Streptococcus mutans*, and *Escherichia coli*; as positive controls and (2) negative controls consisting of sterile saline.

### Covariates

Socio-demographic data on education, income, insurance use, Medicaid status, marital status, parity, maternal age and race and ethnicity were available from maternal self-report. Data on medication use including antibiotics, health behaviors (smoking, alcohol use), and body mass index (BMI); 24-hour dietary recalls during mid-late pregnancy diet were collected over the telephone by a trained nutritionist using the United States Department of Agriculture’s (USDA’s) automated multiple pass method from which we derived the Healthy Eating Index, a measure of dietary quality that compares dietary intake to recommendations from the Dietary Guidelines for Americans^48-50^; clinical conditions (e.g., preeclampsia) were derived from self-report and medical record data.

### Community analysis and statistics

Raw data from the Illumina MiSeq was first converted into FASTQ format 2x312 paired end sequence files using the bcl2fastq program, version 1.8.4, provided by Illumina. Reads were multiplexed using a configuration described previously ^47^. Briefly, for both reads in a pair, the first 24-29 bases were an adapter, followed by an 8-base barcode and an overlapping tag sequence which was followed by a heterogeneity spacer, then a gene specific primer, followed by the target 16S rRNA sequence. Demultiplexed reads were imported into QIIME 2 ^51^, which was used to perform all subsequent processing. Reads were demultiplexed requiring exact barcode matches, and 16S primers were removed allowing 20% mismatches and requiring at least 18 bases. Cleaning, joining, and denoising were performed using DADA2: forward reads were truncated to 275 bps and reverse reads to 260 bps, error profiles were learned with a sample of one million reads, and a maximum expected error of two was allowed. Taxonomic classification was performed with a custom naïve Bayesian classifier trained on the August, 2013 release of GreenGenes^52,53^ and SILVA ^54^. Amplicon sequence variants (ASVs) that could not be classified at least at the phylum level were discarded.

### Statistical Analysis

Descriptive statistics were used to summarize the characteristics of the study population. Means and standard deviations were used for continuous data, and frequencies and percentages for categorical data. Clustering analysis based on the Bray-Curtis dissimilarity measure and the Ward’s linkage was used to construct CSTs. The optimal number of CSTs was determined using silhouette analysis. For alpha diversity, the Shannon index was used to capture the richness and evenness for each sample. Difference in alpha diversity among CSTs was assessed using a regression model with CSTs as the main factor of interest and antibiotic usage, BMI, and race as *a priori* covariates; other covariates were included if they were significant predictors in the model. A mixed-effects model was used to determine an association between EPDS (or PSWQ) and CST (or each taxon) with EPDS (or PSWQ) as the outcome variable, CST (or each taxon) as the variable of interest, and trimester as the random factor. In each model, the covariates (e.g., antibiotic, BMI, race) and an interaction term between CST (or each taxon) and trimester were included. Similar analysis was used to assess an association between each of the sex steroids and CST (or each taxon). P values in univariate analyses were adjusted for multiple testing using the Benjamini-Hochberg (BH) procedure to control false discovery rate (FDR). PICRUSt2^55^, which predict the functional potential of a bacterial community based on marker gene sequencing profiles, was used to predict potential metabolic pathways associated with CST. The point-biserial correlation was used to assess correlations between individual pathways and the presence/absence of individual taxa. All analyses were performed in R.

## RESULTS

### Identification of maternal vaginal microbiome community state types (CST)

The observational cohort of 275 mother-infant dyads yielded 216 third-trimester vaginal samples. Composition of the vaginal microbiota community was quantified by 16S rRNA amplicon sequencing, with taxonomic data obtained from 202 subjects who had a total count larger than 1000 in the taxonomic profile. Amplicon sequence variants (ASVs) at the species level were aggregated at the genus level if their presence was smaller than 10% of samples. The ASVs present at the genus level in less than 10% of samples were removed to reduce potential spurious correlations due to excess zeros, leaving 27 taxa for downstream analysis. These taxa include *Lactobacillus iners, Lactobacillus gasseri*, and *Lactobacillus jensenii*, which were the key vaginal microbiota phylotypes defined by Ravel et al. based on operational taxonomic units (OTUs)^56-58^. Classification of the samples into CSTs provides an analytically tractable summary of the vaginal microbial community, with identification of functionally distinct CSTs that have potential roles in onset of affective symptoms. With a hierarchical clustering analysis based on Bray-Curtis dissimilarity and the silhouette method, we identified three CSTs, dominated by Lactobacillus (CST1), *Lactobacillus iners* (CST2), or no single taxon (CST3) **(Figure 1)**. The lack of a single dominant taxa in CST3 is reflected in a greater Shannon Diversity relative to CST1 and CST2 **(Figure 1)**. In summary, the vaginal microbiota community composition and major phylotypes in our healthy, normative risk pregnant cohort is consistent with previous studies of the healthy pregnant vaginal microbiome^59,60^.

**Figure 1.**
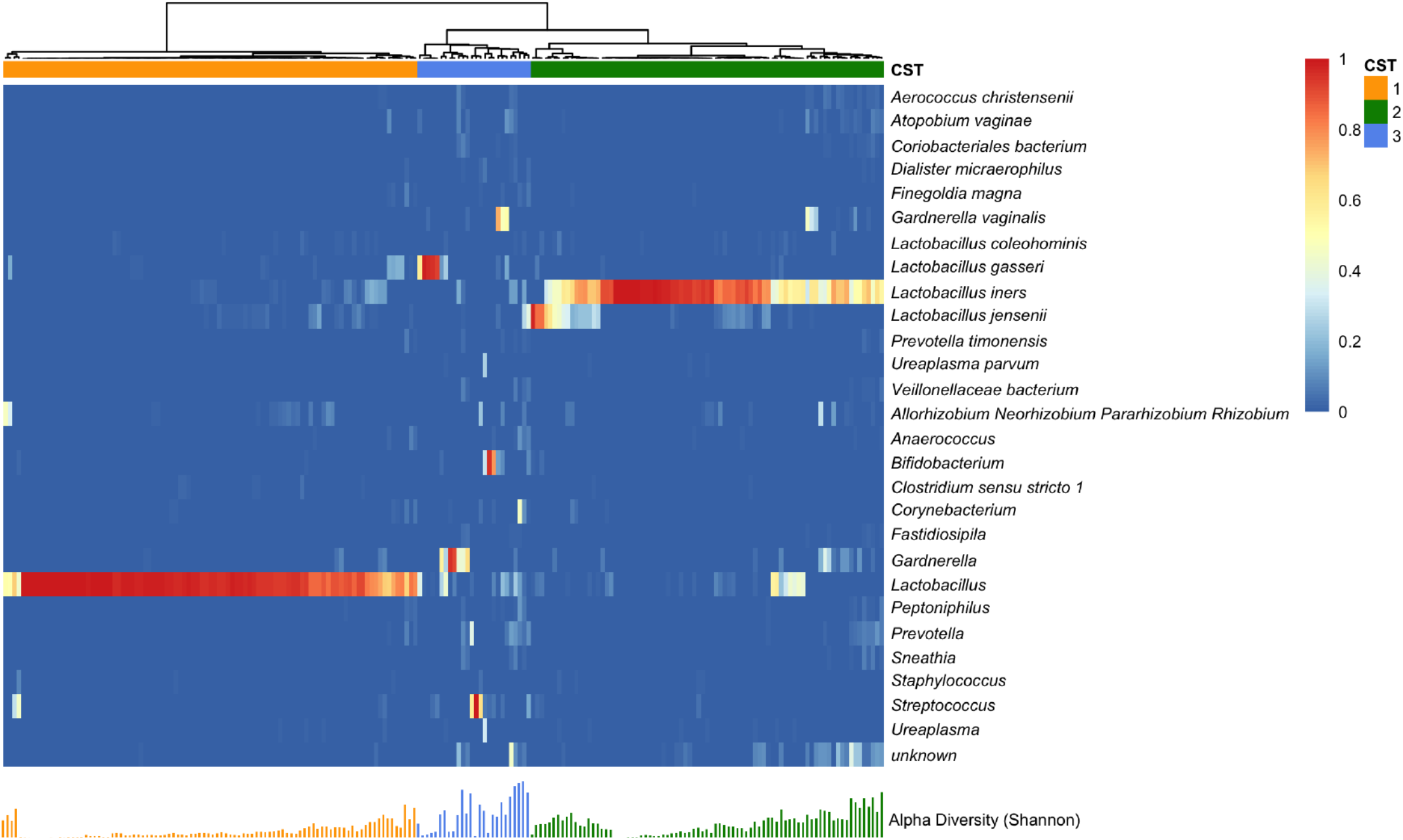
Vaginal Microbiome Profile by Community State Type. Heatmap represents relative abundance of vaginal bacterial taxa (row) in sample (column). Hierarchical clustering with Ward’s linkage clustered samples into three distinct vaginal bacterial compositions called community state type (CST). Each bar under the heatmap indicates Shannon’s index of the corresponding sample.

### Association of vaginal microbiome CSTs and individual taxa with EPDS and PSWQ

To determine potential relationships between affective symptoms and the vaginal microbiota, we applied separate linear mixed effects models to identify statistically significant associations between EPDS and PSWQ scores with CST1-3 and the individual taxa in each CST. For both models, BMI, antibiotic status, and ethnicity/race, and CST or individual taxa were included as fixed effects. We found a statistically significant difference in EPDS scores between CST 1 and CST 3 (*p* = 0.026 at trimester 1; *p* = 0.084 over trimesters 1-3). We also found a statistically significant decrease in PSWQ scores between both CST 2 (*p* = 0.031 at trimester 1; *p* = 0.049 over trimesters 1-3) and CST 3 (*p* = 0.009 at trimester 1; *p* = 0.050 over trimesters 1-3) relative to CST 1 **(Figure 2)**. We used the same analysis strategy for each taxon, with the abundance of the taxon as fixed effects, an interaction term between the taxon abundance and trimester, and trimester as a random effect. With this strategy, we identified 11 taxa associated with PSWQ and EPSD scores (FDR adjusted p-value or *q* ≤ 0.2 at trimester 1). Of these 11 taxa, we found three taxa at *q* = 0.05; *Anaerococcus* (*q* = 0.032 at trimester 1; *q* = 0.101 over trimesters 1-3), *Lactobacillus* (*q* = 0.032 at trimester 1; *q* = 0.101 over trimesters 1-3), and *Peptoniphilus* (*q* = 0.046 at trimester 1; *q* = 0.186 over trimesters 1-3) with PSWQ score **(Figure 3)**. Overall, relative to CST1, CST3 was more abundant in subjects with lower EPDS scores, and both CST2 and CST3 were more abundant in those with lower PSWQ. At the genus level, significant negative associations were identified with *Anaerococcus, Peptoniphilus*, and PSWQ. In contrast, a significant positive association was identified between *Lactobacillus* and PSWQ.

**Figure 2.**
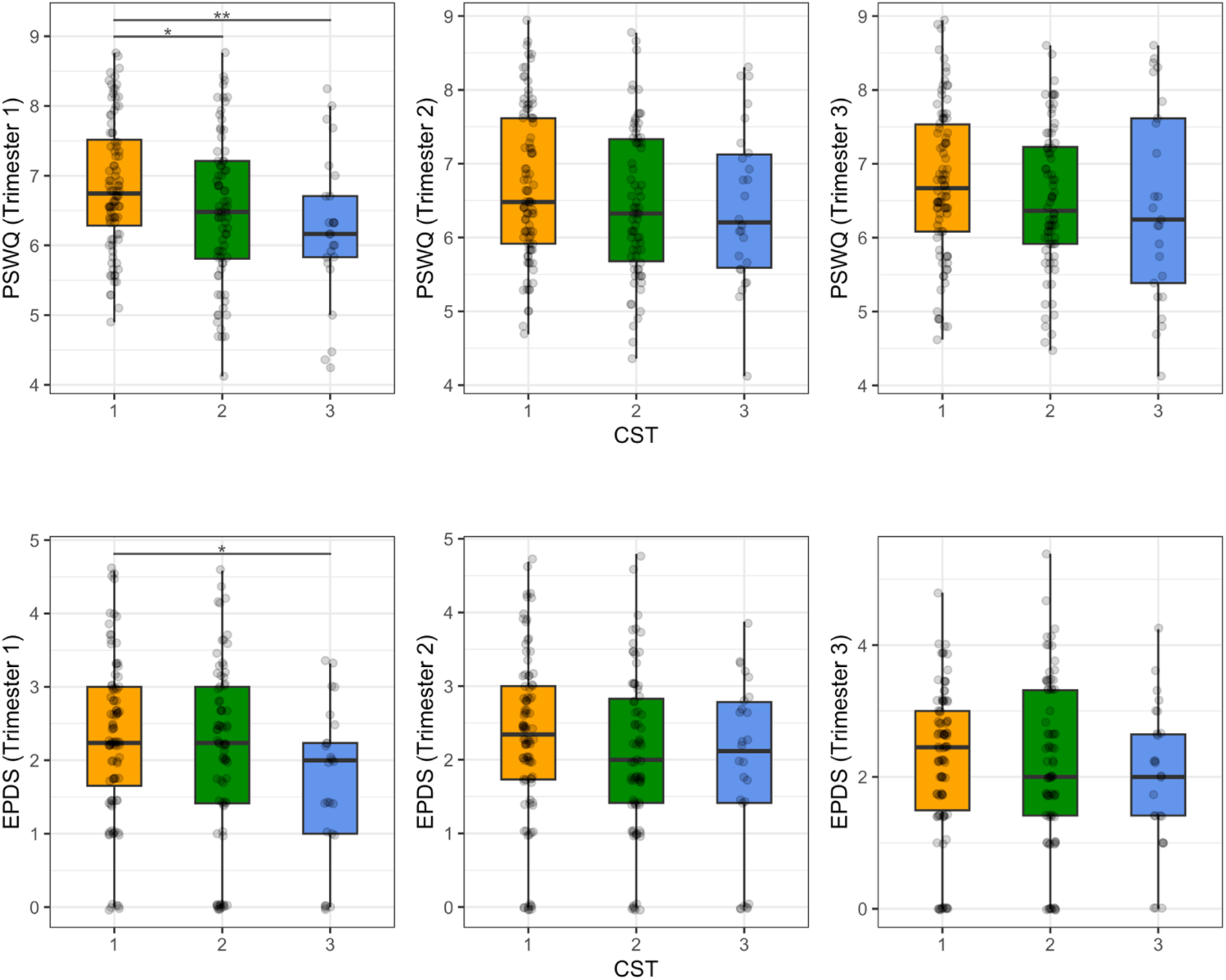
Distribution of PSWQ/EPDS Scores for CST at each trimester. * denotes *p* ≤ 0.05 and ** denotes *p* ≤ 0.01.

**Figure 3.**
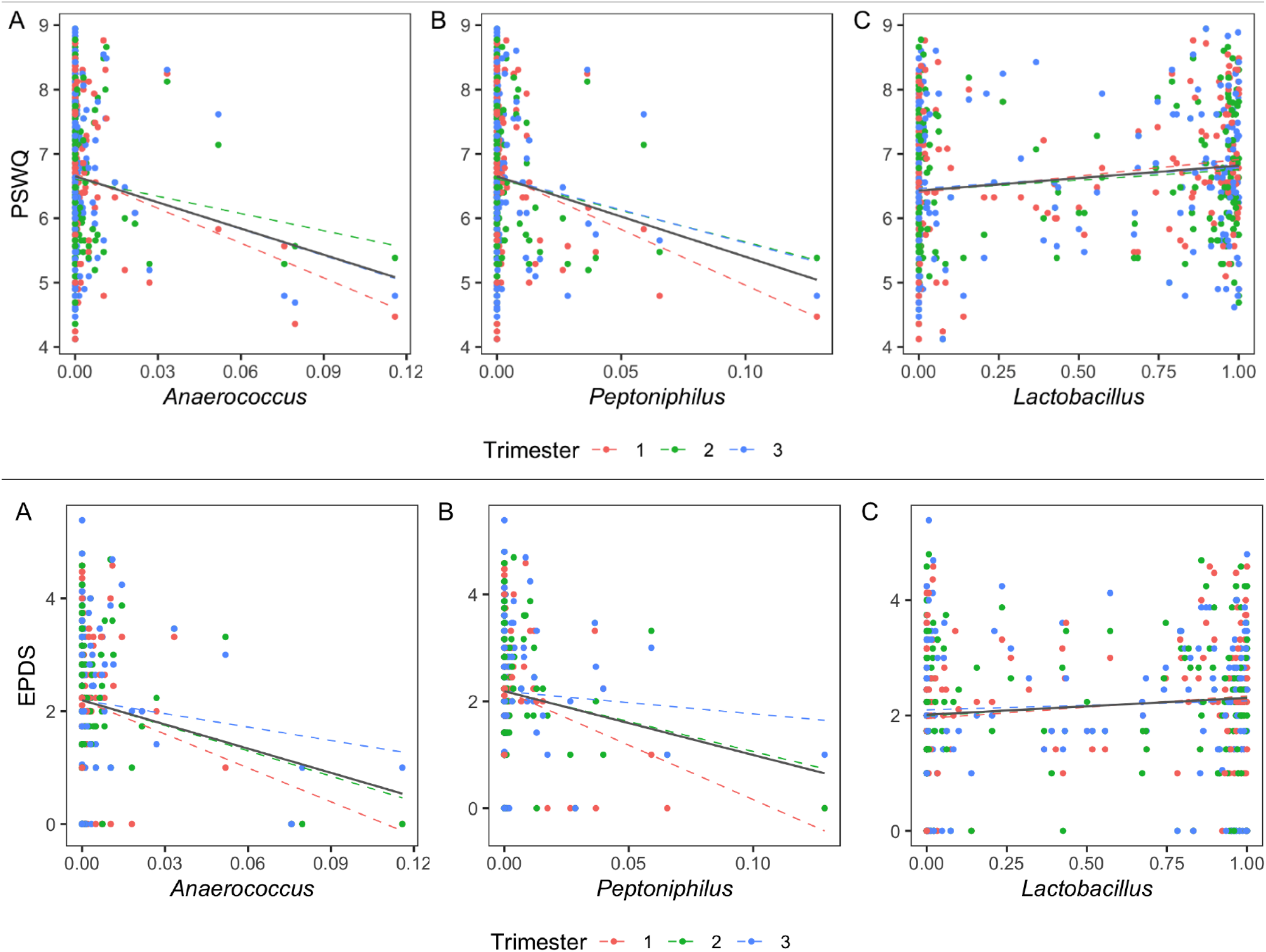
Correlation Between Bacterial Taxa and PSWQ/EPDS. Scatter plots show the association between PSWQ/EPDS scores and the relative abundance of taxa at q = 0.05. Each dotted color line represents a linear relation between PSWQ/EPDS and each taxon at each trimester. The solid black line represents an overall linear relation between PSWQ/EPDS and each taxon across trimesters.

### Microbiome metabolic pathways associated with CST, individual taxa, EPDS, and PSWQ

To determine potential associations between vaginal microbiota function, CST, and individual taxa, we used PICRUSt2^55^ to identify predicted metabolic pathways in the individual vaginal samples and differential abundance of these pathways in CST1-3 and the 11 individual taxa in these CSTs associated with PSWQ and/or EPDS. To determine differentially abundant pathways between CSTs, we first filtered out pathways that appeared in fewer than 10% of samples or had a maximum relative abundance smaller than 0.001. We then applied the centered log-ratio transformation after replacing zeros with 0.5 and performed a Kruskal-Wallis test for each pathway. After multiple comparison correction using the Benjamini-Hochberg method to control false discovery rate (FDR), we identified 76 pathways with *q* ≤ 0.05 **(Figure 4)**. A full list of identified pathways can be found in **Supplemental Table 1**. The pathways for L-tryptophan and L-phenylalanine; biosynthetic precursors of serotonin, dopamine and norepinephrine, have a significant positive association with CST3 (*q* = 6.86 × 10 ^−8^ for L-phenylalanine and *q* = 1.41 × 10^−3^ for L-tryptophan), a CST associated with improved EPDS and PSWQ scores **(Figure 3)**. To assess the association between these two pathways and the 11 taxa associated with maternal anxiety and depression (MIC in Figure 3), we used the point-biserial correlation, i.e., correlation between the abundance of a pathway and the presence/absence of a taxon, to alleviate the spurious correlation due to excess zeros in the taxonomic profile. We identified significant correlations of the L-tryptophan pathway with *Finegoldia magna* (*r* = 0.097), *Coriobaceriales* (*r* = 0.099), *Fastidiosipila* (*r* = 0.067), and *Peptoniphilus* (*r* = 0.077) and the L-phenylalanine pathway with *Fingoldia magna* (*r* = 0.097), *Prevotella timonensis* (*r* = 0.153), *Peptoniphilus* (*r* = 0.136), and *Lactobacillus* (*r* = 0.105) **(Figure 4)**. These results suggest the metabolic pathways known to interact with psychosocial stress may be differentially expressed in pregnant people with affective symptoms.

**Figure 4.**
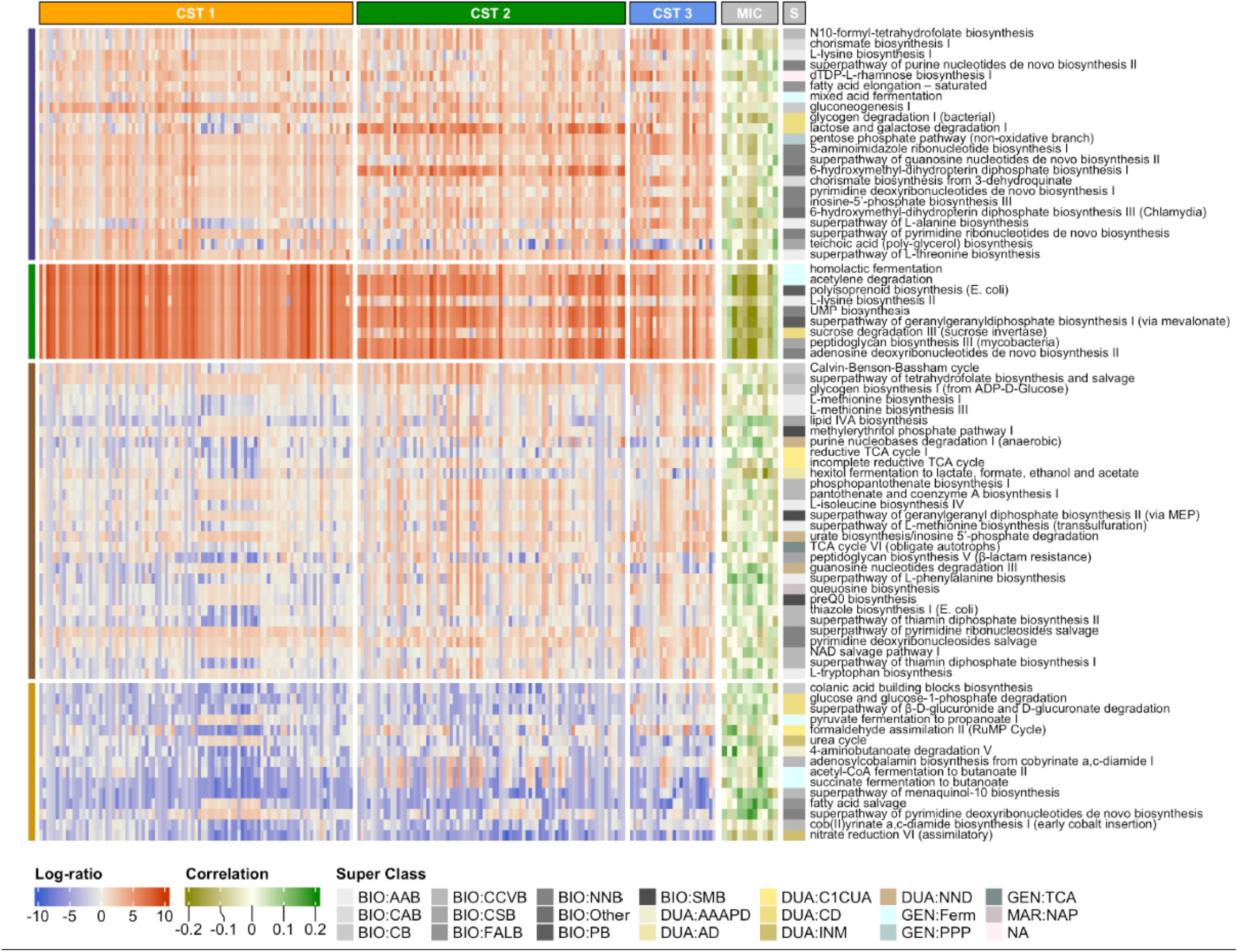
Association of Predicted Metabolic Pathways with Community State Type, Individual Taxa, A subset of pathways significantly associated with the community state type and their correlations with taxa related to maternal anxiety/depression. Column annotation CST indicates the community state type, MIC indicates taxa, and S indicates the super class of each pathway. The taxa are, from the left to the right, *Anaerococcus, Lactobacillus, Peptoniphilus, Veillonellaceae bacterium, Fastidiosipila, Coriobacteriales bacterium, Sneathia, Prevotella timonensis, Prevotella, Finegoldia magna*, and *Corynebacterium*. The full pathway names for the Super Class are described in **Supplementary Table 1**.

### Association of CSTs with biomarkers of affective symptoms and stress

Given previous research that cortisol and sex steroids may be biomarkers of affective symptoms and stress, we hypothesized that these antenatal hormones would also be associated with maternal vaginal microbiota. In the subset of subjects included in this analysis, cortisol diurnal slope was associated with EPDS (coefficient = 0.859, *p* = 0.011), but not PSWQ. Sex steroid levels (testosterone (T), Estrone (E1), Estradiol (E2), and Estriol (E3)) were not associated with either EPDS or PSWQ **(Table 1)**. We next examined associations between CST1-3, individual taxa, sex steroids and cortisol. We fit separate linear mixed effects models with each sex steroid level as the response, and the same fixed and random effects that we used for looking at associations between depression and anxiety scores and CSTs. A statistically significant association was determined between CST1 and CST3 for free or bioavailable testosterone (Free T Ng/Dl (*p* = 0.025)); none of the other sex steroid levels had a statistically significant association with the CSTs. We did not identify significant associations between the sex steroid levels and individual taxa. Cortisol was not significantly associated with either CST or individual taxa. Together these data points suggest that the hormonal physiology of affective symptoms may play less of a role in shaping the vaginal microbiome during pregnancy.

**Table 1.** Mother’s Demographics and Clinical Characteristics and Their Associations with EPDS and PSWQ.

|  | Summary | EPDS Coefficient (SE) | PSWQ Coefficient (SE) |
| --- | --- | --- | --- |
| Maternal BMI | 27.8 (6.8) | 0.037 (0.048); p=0.446 | -0.054 (0.134); p=0.686 |
| Race/Ethnicity (n=202) | White: 0.604 (122)<br>Black: 0.228 (46)<br>Other: 0.168 (34) | 1.727 (0.792); p=0.030<br>2.011 (0.880); p=0.023 | -1.775 (2.243); p=0.430<br>2.757 (2.494); p=0.270 |
| Maternal Smoking (n=202) | No: 0.933 (181)<br>Yes: 0.067 (13) | 1.363 (1.051); p=0.195 | -3.039 (2.773); p=0.274 |
| Parity (n=202) | No: 0.302 (61)<br>Yes: 0.698 (141) | -0.359 (0.710); p=0.613 | -1.586 (1.982); p=0.430 |
| Antibiotics within 4 weeks prior to sampling (n=202) | No: 0.969 (6)<br>Yes: 0.031 (189) | -0.188 (0.801); p=0.815 | -1.257 (1.850); p=0.497 |
| Fetal sex (n=202) | M: 0.515 (104)<br>F: 0.485 (98) | -0.070 (0.649); p=0.914 | -2.020 (1.810); p=0.266 |
| GDM.Final (n=202) | No: 0.96 (194)<br>Yes: 0.04 (8) | -0.574 (1.654); p=0.729 | -1.140 (4.628); p=0.806 |
| Maternal Age | 28.9 (9.6) | -0.097 (0.071); p=0.174 | -0.240 (0.199); p=0.229 |
| Cortisol (Diurnal Slope) | -0.6 (0.5) | 0.859 (0.336); p=0.011 | 0.771 (0.832); p=0.355 |
| Cortisol (AUC) | 76.8 (39.4) | 0.002 (0.004); p=0.578 | 0.001 (0.011); p=0.908 |
| Total.Free.T.Percent | 0.524 (0.148) | 0.623 (1.100); p=0.571 | 1.397 (2.625); p=0.594 |
| E1.Pg.MI $\times 10^{-3}$ | 3.886 (3.884) | 0.013 (0.041); p=0.749 | -0.110 (0.103); p=0.285 |
| E2.Pg.MI $\times 10^{-3}$ | 6.517 (5.018) | 0.046 (0.027); p=0.096 | -0.060 (0.070); p=0.390 |
| E3.Pg.MI $\times 10^{-3}$ | 3.560 (3.355) | 0.006 (0.040); p=0.889 | -0.153 (0.099); p=0.122 |

The table above shows summary statistics and the associations between depression and anxiety scores for various demographic variables. For continuous variables, the summary statistics are reported as mean (standard deviation). For binary variables, the summary statistics are reported as the sample proportion of those who have that characteristic and the total number of responses. For the EPDS and PSWQ columns, we have reported the regression coefficients and their corresponding standard error and p-value when fitting a linear mixed effects model using EPDS/PSWQ as the response variable.

## DISCUSSION

The current study supports the hypothesis that affective symptoms associate with the microbiota, and further advances this field in several critical directions. First, we included multiple measures of affective distress measured across gestation, including affective symptoms of depression, anxiety, and life event stress. In addition to testing the robustness of association across measures of distress at multiple times during pregnancy, we also include detailed assessments of possible confounding factors, including socio-demographic^61^, diet, and medications. We identified three dominant *Lactobacillus* phylotypes, *L. iners, L. gasseri*, and *L. jensenii*, in the third trimester maternal vaginal microbiome. Second, we assessed microbiome composition and clustered the taxa into Community State Types (CSTs), which collapse the variation in microbiota composition into archetypal states that represent of the microbiota and its functional signature. Individual taxa clustered into three major CSTs (CST1-3) based on their vaginal microbiota compositional dissimilarity **(Figure 1)**, with CST1 dominated by Lactobacillus, CST2 dominated by Lactobacillus inners, and CST3 not dominated by a single taxon. Linear mixed effects models with individual taxa revealed an association with increased abundance of *Peptoniphilus, Anaerococcus*, and *Lactobacillus* and improved EPDS and PSWQ scores **(Figure 2)**. Similar results were observed in a previous study on gut microbiota and metabolites on prenatal depression, with greater abundance of *Peptoniphilus* and *Anaerococcus* associated with diminished prenatal depression^62^.

We further extended the association of individual taxa and CSTs with affective symptoms and predicted biological pathway abundance derived from 16S rRNA taxa. Our prediction of functional pathway abundances^55^ **(Figure 3)** identified significant correlations of multiple taxon with the L-tryptophan and L-phenylalanine functional pathways, the precursors of serotonin, dopamine, and norepinephrine -- key neurotransmitters associated with depression^63-65^. A single taxon, *Peptoniphilus*, was significantly correlated with both functional pathways. Linear mixed models of CSTs revealed a significant association with CST3 and lower EPDS and PSWQ scores. Notably, *Peptoniphilus* was associated with lower EPDS and PSWQ both as an individual taxon and as a member of CST3, revealing a potential key role of this bacterium in affective disorders.

A particularly novel extension of this work is the consideration of biological markers of affective distress that may provide a mechanistic link with CSTs. Glucocorticoids, indexed by diurnal salivary cortisol, are associated with affective symptoms, and may account for their effects on maternal-fetal-placenta health; we examine their association with vaginal microbiome CSTs. A second biological marker of affective distress that may explain a link with the vaginal microbiome are sex steroids. Estrogen-based biological pathways are of special interest with indirect support from studies which suggest that estrogen-containing contraceptives are associated with lower risk of bacterial vaginosis^66^. We did not detect reliable associations between vaginal microbiome and maternal cortisol or estrogens, suggesting that these markers do not mediate the link between clinical measures and vaginal microbiome - and neither do they have a link independent of clinical symptoms and stress. Previous studies reporting associations between biomarkers of symptom or stress and prenatal maternal microbiome have been based on animals, gut microbiome, or measures of diversity^67^. The observed association between maternal prenatal testosterone and vaginal microbiome was unanticipated and requires further study and replication. As a result, although there is accumulating evidence, including from the current paper, that clinical measures of distress are associated with maternal microbiome composition, the mechanisms underlying this connection remain unclear.

The study has several limitations. One is that our findings do not discriminate between two potential alternatives; that affective symptoms shape the maternal vaginal microbiome or that alteration of the vaginal microbiota influence onset of affective symptoms. Longitudinal collection of each is needed to differentiate these accounts, and pre-pregnancy vaginal microbiome would be especially value - although challenging to obtain. A second limitation is that we focused on 16S rRNA; further metagenomics analysis is needed to confirm the biological pathways suggested in this paper. Additionally, the findings obtained here were derived from a generally health, diverse sample and may not generalize to other populations. Finally, the current study focused on only one maternal microbiome and so we are unable to determine how specific the findings obtained here are particular to the vaginal microbiome and not also found with another maternal microbiota, e.g., gut microbiome. Set against these limitations are several strengths, including detailed assessment of symptoms and covariates in well-characterized samples.

The findings suggest several future directions. Perhaps most importantly, these results provide a basis for understanding one route by which prenatal affective distress may alter child health outcomes. Further research to test that hypothesis is needed, as the early seeding of the infant microbiome from maternal (vaginal) microbiome may shape child immune and neurodevelopmental health outcomes. The clinical implications of the work are not yet clear and require replication. It remains an important possibility that the maternal prenatal microbiomes could provide intervention target for promoting maternal, perinatal, and child health outcomes.

## Availability of Data and Materials

Illumina 16S rRNA V3V4 amplicons were deposited in Sequence Read Archive under BioProject PRJNA1099167, including positive and negative controls on each plate.

## Author Contributions

KS, TGO, and SRG designed the study and completed the initial drafts of the manuscript. MS, RB, and XQ carried out biostatistical analyses. ALG carried out 16S rRNA microbiome sequencing and analyses. JNM contributed to microbiome analyses. JB carried out collection of socio-demographic and clinical data from the mother-infant cohort. RM, EB, MS, RB, KS, TGO and SRG were responsible for final revisions of the manuscript. All authors read the final draft of the manuscript.

## Acknowledgements

We thank the Genomics Research Center for performing 16S rRNA sequencing.

## REFERENCES

1. Capron LE, Glover V, Pearson RM, et al. Associations of maternal and paternal antenatal mood with offspring anxiety disorder at age 18 years. J Affect Disord. Nov 15 2015;187:20–6. doi:10.1016/j.jad.2015.08.012

2. Grote NK, Bridge JA, Gavin AR, Melville JL, Iyengar S, Katon WJ. A meta-analysis of depression during pregnancy and the risk of preterm birth, low birth weight, and intrauterine growth restriction. Arch Gen Psychiatry. Oct 2010;67(10):1012–24. doi:10.1001/archgenpsychiatry.2010.111

3. Raina J, El-Messidi A, Badeghiesh A, Tulandi T, Nguyen TV, Suarthana E. Pregnancy hypertension and its association with maternal anxiety and mood disorders: A population-based study of 9 million pregnancies. J Affect Disord. Feb 15 2021;281:533–538. doi:10.1016/j.jad.2020.10.058

4. Hechler C, Borewicz K, Beijers R, et al. Association between Psychosocial Stress and Fecal Microbiota in Pregnant Women. Sci Rep. Mar 14 2019;9(1):4463. doi:10.1038/s41598-019-40434-8

5. Radjabzadeh D, Bosch JA, Uitterlinden AG, et al. Gut microbiome-wide association study of depressive symptoms. Nat Commun. Dec 6 2022;13(1):7128. doi:10.1038/s41467-022-34502-3

6. Jasarevic E, Howard CD, Morrison K, et al. The maternal vaginal microbiome partially mediates the effects of prenatal stress on offspring gut and hypothalamus. Nat Neurosci. Aug 2018;21(8):1061–1071. doi:10.1038/s41593-018-0182-5

7. Jasarevic E, Howerton CL, Howard CD, Bale TL. Alterations in the Vaginal Microbiome by Maternal Stress Are Associated With Metabolic Reprogramming of the Offspring Gut and Brain. Endocrinology. Sep 2015;156(9):3265–76. doi:10.1210/en.2015-1177

8. Borgogna JC, Anastario M, Firemoon P, et al. Vaginal microbiota of American Indian women and associations with measures of psychosocial stress. PLoS One. 2021;16(12):e0260813. doi:10.1371/journal.pone.0260813

9. Gerson KD, Liao J, McCarthy C, et al. A non-optimal cervicovaginal microbiota in pregnancy is associated with a distinct metabolomic signature among non-Hispanic Black individuals. Sci Rep. Nov 23 2021;11(1):22794. doi:10.1038/s41598-021-02304-0

10. Gerson KD, McCarthy C, Ravel J, Elovitz MA, Burris HH. Effect of a Nonoptimal Cervicovaginal Microbiota and Psychosocial Stress on Recurrent Spontaneous Preterm Birth. Am J Perinatol. Apr 2021;38(5):407–413. doi:10.1055/s-0040-1717098

11. Turpin R, Slopen N, Borgogna JC, et al. Perceived Stress and Molecular Bacterial Vaginosis in the National Institutes of Health Longitudinal Study of Vaginal Flora. Am J Epidemiol. Nov 2 2021;190(11):2374–2383. doi:10.1093/aje/kwab147

12. Romero R, Hassan SS, Gajer P, et al. The vaginal microbiota of pregnant women who subsequently have spontxaneous preterm labor and delivery and those with a normal delivery at term. Microbiome. 2014;2:18. doi:10.1186/2049-2618-2-18

13. Cryan JF, Dinan TG. Mind-altering microorganisms: the impact of the gut microbiota on brain and behaviour. Nat Rev Neurosci. Oct 2012;13(10):701–12. doi:10.1038/nrn3346

14. Golubeva AV, Crampton S, Desbonnet L, et al. Prenatal stress-induced alterations in major physiological systems correlate with gut microbiota composition in adulthood. Psychoneuroendocrinology. Oct 2015;60:58–74. doi:10.1016/j.psyneuen.2015.06.002

15. Gur TL, Shay L, Palkar AV, et al. Prenatal stress affects placental cytokines and neurotrophins, commensal microbes, and anxiety-like behavior in adult female offspring. Brain Behav Immun. Aug 2017;64:50–58. doi:10.1016/j.bbi.2016.12.021

16. Jasarevic E, Hill EM, Kane PJ, et al. The composition of human vaginal microbiota transferred at birth affects offspring health in a mouse model. Nat Commun. Nov 1 2021;12(1):6289. doi:10.1038/s41467-021-26634-9

17. Rifkin-Graboi A, Meaney MJ, Chen H, et al. Antenatal maternal anxiety predicts variations in neural structures implicated in anxiety disorders in newborns. J Am Acad Child Adolesc Psychiatry. Apr 2015;54(4):313–21 e2. doi:10.1016/j.jaac.2015.01.013

18. Sampson TR, Mazmanian SK. Control of brain development, function, and behavior by the microbiome. Cell Host Microbe. May 13 2015;17(5):565–76. doi:10.1016/j.chom.2015.04.011

19. Zijlmans MA, Korpela K, Riksen-Walraven JM, de Vos WM, de Weerth C. Maternal prenatal stress is associated with the infant intestinal microbiota. Psychoneuroendocrinology. Mar 2015;53:233–45. doi:10.1016/j.psyneuen.2015.01.006

20. Hourigan SK, Dominguez-Bello MG, Mueller NT. Can maternal-child microbial seeding interventions improve the health of infants delivered by Cesarean section? Cell Host Microbe. May 11 2022;30(5):607–611. doi:10.1016/j.chom.2022.02.014

21. Song SJ, Wang J, Martino C, et al. Naturalization of the microbiota developmental trajectory of Cesarean-born neonates after vaginal seeding. Med. Aug 13 2021;2(8):951–964 e5. doi:10.1016/j.medj.2021.05.003

22. Vatanen T, Jabbar KS, Ruohtula T, et al. Mobile genetic elements from the maternal microbiome shape infant gut microbial assembly and metabolism. Cell. Dec 22 2022;185(26):4921–4936 e15. doi:10.1016/j.cell.2022.11.023

23. Bogaert D, van Beveren GJ, de Koff EM, et al. Mother-to-infant microbiota transmission and infant microbiota development across multiple body sites. Cell Host Microbe. Mar 8 2023;31(3):447–460 e6. doi:10.1016/j.chom.2023.01.018

24. Brodin P. Immune-microbe interactions early in life: A determinant of health and disease long term. Science. May 27 2022;376(6596):945–950. doi:10.1126/science.abk2189

25. Cox LM, Yamanishi S, Sohn J, et al. Altering the intestinal microbiota during a critical developmental window has lasting metabolic consequences. Cell. Aug 14 2014;158(4):705–721. doi:10.1016/j.cell.2014.05.052

26. Dominguez-Bello MG, Costello EK, Contreras M, et al. Delivery mode shapes the acquisition and structure of the initial microbiota across multiple body habitats in newborns. Research Support, N.I.H., Extramural Research Support, Non-U.S. Gov’t. Proc Natl Acad Sci U S A. Jun 29 2010;107(26):11971–5. doi:10.1073/pnas.1002601107

27. Jasarevic E, Bale TL. Prenatal and postnatal contributions of the maternal microbiome on offspring programming. Front Neuroendocrinol. Oct 2019;55:100797. doi:10.1016/j.yfrne.2019.100797

28. Galley JD, Mashburn-Warren L, Blalock LC, et al. Maternal anxiety, depression and stress affects offspring gut microbiome diversity and bifidobacterial abundances. Brain Behav Immun. Jan 2023;107:253–264. doi:10.1016/j.bbi.2022.10.005

29. Cryan JF, O’Riordan KJ, Cowan CSM, et al. The Microbiota-Gut-Brain Axis. Physiol Rev. Oct 1 2019;99(4):1877–2013. doi:10.1152/physrev.00018.2018

30. Carlson AL, Xia K, Azcarate-Peril MA, et al. Infant Gut Microbiome Associated With Cognitive Development. Biol Psychiatry. Jan 15 2018;83(2):148–159. doi:10.1016/j.biopsych.2017.06.021

31. Carlson AL, Xia K, Azcarate-Peril MA, et al. Infant gut microbiome composition is associated with non-social fear behavior in a pilot study. Nat Commun. Jun 2 2021;12(1):3294. doi:10.1038/s41467-021-23281-y

32. Cowan CSM, Dinan TG, Cryan JF. Annual Research Review: Critical windows - the microbiota-gut-brain axis in neurocognitive development. J Child Psychol Psychiatry. Nov 26 2019;doi:10.1111/jcpp.13156

33. DiGiulio DB, Callahan BJ, McMurdie PJ, et al. Temporal and spatial variation of the human microbiota during pregnancy. Proc Natl Acad Sci U S A. Sep 1 2015;112(35):11060–5. doi:10.1073/pnas.1502875112

34. Hu J, Ly J, Zhang W, et al. Microbiota of newborn meconium is associated with maternal anxiety experienced during pregnancy. Dev Psychobiol. Jul 2019;61(5):640–649. doi:10.1002/dev.21837

35. Warner BB. The contribution of the gut microbiome to neurodevelopment and neuropsychiatric disorders. Pediatr Res. Jan 2019;85(2):216–224. doi:10.1038/s41390-018-0191-9

36. O’Connor T, Best M, Brunner J, et al. Cohort profile: Understanding Pregnancy Signals and Infant Development (UPSIDE): a pregnancy cohort study on prenatal exposure mechanisms for child health. BMJ Open. Apr 1 2021;11(4):e044798. doi:10.1136/bmjopen-2020-044798

37. Knapp EA, Kress AM, Parker CB, et al. The Environmental Influences on Child Health Outcomes (ECHO)-Wide Cohort. Am J Epidemiol. Aug 4 2023;192(8):1249–1263. doi:10.1093/aje/kwad071

38. Cox JL, Holden JM, Sagovsky R. Detection of postnatal depression. Development of the 10-item Edinburgh Postnatal Depression Scale. Br J Psychiatry. Jun 1987;150:782–6.

39. Murray L, Carothers AD. The validation of the Edinburgh Post-natal Depression Scale on a community sample. Br J Psychiatry. Aug 1990;157:288–90.

40. Barnett BE, Hanna B, Parker G. Life event scales for obstetric groups. J Psychosom Res. 1983;27(4):313–20. doi:10.1016/0022-3999(83)90054-5

41. Hansel MC, Murphy HR, Brunner J, et al. Associations between neighborhood stress and maternal sex steroid hormones in pregnancy. BMC Pregnancy Childbirth. Oct 16 2023;23(1):730. doi:10.1186/s12884-023-06043-0

42. Shipp GM, Wosu AC, Knapp EA, et al. Maternal Pre-Pregnancy BMI, Breastfeeding, and Child BMI. Pediatrics. Jan 1 2024;153(1)doi:10.1542/peds.2023-061466

43. http://www.macses.ucsf.edu/research/allostatic/notebook/salivarycort.html.

44. Pruessner JC, Kirschbaum C, Meinlschmid G, Hellhammer DH. Two formulas for computation of the area under the curve represent measures of total hormone concentration versus time-dependent change. Psychoneuroendocrinology. Oct 2003;28(7):916–31. doi:S0306453002001087 [pii]

45. Shiraishi S, Lee PW, Leung A, Goh VH, Swerdloff RS, Wang C. Simultaneous measurement of serum testosterone and dihydrotestosterone by liquid chromatography-tandem mass spectrometry. Clin Chem. Nov 2008;54(11):1855–63. doi:10.1373/clinchem.2008.103846

46. Qoubaitary A, Meriggiola C, Ng CM, et al. Pharmacokinetics of testosterone undecanoate injected alone or in combination with norethisterone enanthate in healthy men. J Androl. Nov-Dec 2006;27(6):853–67. doi:10.2164/jandrol.106.000281

47. Holm JB, Humphrys MS, Robinson CK, et al. Ultrahigh-Throughput Multiplexing and Sequencing of >500-Base-Pair Amplicon Regions on the Illumina HiSeq 2500 Platform. mSystems. Jan-Feb 2019;4(1)doi:10.1128/mSystems.00029-19

48. Reedy J, Lerman JL, Krebs-Smith SM, et al. Evaluation of the Healthy Eating Index-2015. J Acad Nutr Diet. Sep 2018;118(9):1622–1633. doi:10.1016/j.jand.2018.05.019

49. Reedy J, Subar AF. 90th Anniversary Commentary: Diet Quality Indexes in Nutritional Epidemiology Inform Dietary Guidance and Public Health. J Nutr. Oct 1 2018;148(10):1695–1697. doi:10.1093/jn/nxy184

50. Reedy J, Subar AF, George SM, Krebs-Smith SM. Extending Methods in Dietary Patterns Research. Nutrients. May 7 2018;10(5)doi:10.3390/nu10050571

51. Bolyen E, Rideout JR, Dillon MR, et al. Reproducible, interactive, scalable and extensible microbiome data science using QIIME 2. Nat Biotechnol. Aug 2019;37(8):852–857. doi:10.1038/s41587-019-0209-9

52. DeSantis TZ, Hugenholtz P, Larsen N, et al. Greengenes, a chimera-checked 16S rRNA gene database and workbench compatible with ARB. Appl Environ Microbiol. Jul 2006;72(7):5069–72. doi:10.1128/AEM.03006-05

53. McDonald D, Price MN, Goodrich J, et al. An improved Greengenes taxonomy with explicit ranks for ecological and evolutionary analyses of bacteria and archaea. ISME J. Mar 2012;6(3):610–8. doi:10.1038/ismej.2011.139

54. Quast C, Pruesse E, Yilmaz P, et al. The SILVA ribosomal RNA gene database project: improved data processing and web-based tools. Nucleic Acids Res. Jan 2013;41(Database issue):D590–6. doi:10.1093/nar/gks1219

55. Douglas GM, Maffei VJ, Zaneveld JR, et al. PICRUSt2 for prediction of metagenome functions. Nat Biotechnol. Jun 2020;38(6):685–688. doi:10.1038/s41587-020-0548-6

56. Gajer P, Brotman RM, Bai G, et al. Temporal dynamics of the human vaginal microbiota. Sci Transl Med. May 2 2012;4(132):132ra52. doi:10.1126/scitranslmed.3003605

57. Holm JB, France MT, Gajer P, et al. Integrating compositional and functional content to describe vaginal microbiomes in health and disease. Microbiome. Nov 30 2023;11(1):259. doi:10.1186/s40168-023-01692-x

58. Symul L, Jeganathan P, Costello EK, et al. Sub-communities of the vaginal microbiota in pregnant and non-pregnant women. Proc Biol Sci. Nov 29 2023;290(2011):20231461. doi:10.1098/rspb.2023.1461

59. Baud A, Hillion KH, Plainvert C, et al. Microbial diversity in the vaginal microbiota and its link to pregnancy outcomes. Sci Rep. Jun 4 2023;13(1):9061. doi:10.1038/s41598-023-36126-z

60. Romero R, Hassan SS, Gajer P, et al. The composition and stability of the vaginal microbiota of normal pregnant women is different from that of non-pregnant women. Microbiome. Feb 3 2014;2(1):4. doi:10.1186/2049-2618-2-4

61. Dixon M, Dunlop AL, Corwin EJ, Kramer MR. Joint effects of individual socioeconomic status and residential neighborhood context on vaginal microbiome composition. Front Public Health. 2023;11:1029741. doi:10.3389/fpubh.2023.1029741

62. Xie T, Fan X, Pang H, et al. Association between gut microbiota and its functional metabolites with prenatal depression in women. Neurobiol Stress. Jan 2024;28:100592. doi:10.1016/j.ynstr.2023.100592

63. Correia AS, Vale N. Tryptophan Metabolism in Depression: A Narrative Review with a Focus on Serotonin and Kynurenine Pathways. Int J Mol Sci. Jul 31 2022;23(15)doi:10.3390/ijms23158493

64. Moncrieff J, Cooper RE, Stockmann T, Amendola S, Hengartner MP, Horowitz MA. The serotonin theory of depression: a systematic umbrella review of the evidence. Mol Psychiatry. Aug 2023;28(8):3243–3256. doi:10.1038/s41380-022-01661-0

65. Zhou M, Fan Y, Xu L, et al. Microbiome and tryptophan metabolomics analysis in adolescent depression: roles of the gut microbiota in the regulation of tryptophan-derived neurotransmitters and behaviors in human and mice. Microbiome. Jun 30 2023;11(1):145. doi:10.1186/s40168-023-01589-9

66. Graham ME, Herbert WG, Song SD, et al. Gut and vaginal microbiomes on steroids: implications for women’s health. Trends Endocrinol Metab. Aug 2021;32(8):554–565. doi:10.1016/j.tem.2021.04.014

67. Kimmel MC, Verosky B, Chen HJ, Davis O, Gur TL. The Maternal Microbiome as a Map to Understanding the Impact of Prenatal Stress on Offspring Psychiatric Health. Biol Psychiatry. Feb 15 2024;95(4):300–309. doi:10.1016/j.biopsych.2023.11.014

