## Supplemental Table 1 for "Affective Symptoms in Pregnancy are Associated with the Vaginal Microbiome"

**Supplemental Table 1. Full Pathway Names of Supper Class in Figure 4**

| | |
|---|---|
| BIO:AAB | Biosynthesis → Amino Acid Biosynthesis |
| BIO:CAB | Biosynthesis → Carboxylic Acid Biosynthesis |
| BIO:CB | Biosynthesis → Carbohydrate Biosynthesis |
| BIO:CCVB | Biosynthesis → Cofactor, Carrier, and Vitamin Biosynthesis |
| BIO:CSB | Biosynthesis → Cell Structure Biosynthesis |
| BIO:FALB | Biosynthesis → Fatty Acid and Lipid Biosynthesis |
| BIO:NNB | Biosynthesis → Nucleoside and Nucleotide Biosynthesis |
| BIO:PB | Biosynthesis → Polyprenyl Biosynthesis |
| BIO:SMB | Biosynthesis → Secondary Metabolite Biosynthesis |
| BIO:OTHER | Biosynthesis → Other Biosynthesis |
| DUA:AAAPD | Degradation/Utilization/Assimilation → Amide, Amidine, Amine, and Polyamine Degradation |
| DUA:AD | Degradation/Utilization/Assimilation → Alcohol Degradation |
| DUA:C1CUA | Degradation/Utilization/Assimilation → C1 Compound Utilization and Assimilation |
| DUA:CD | Degradation/Utilization/Assimilation → Carbohydrate Degradation |
| DUA:INM | Degradation/Utilization/Assimilation → Inorganic Nutrient Metabolism |
| DUA:NND | Degradation/Utilization/Assimilation → Nucleoside and Nucleotide Degradation |
| GEN:Ferm | Generation → Fermentation Generation of Precursor Metabolites and Energy |
| GEN:PPP | Generation → Pentose Phosphate Pathways Generation of Precursor Metabolites and Energy |
| GEN:TCA | Generation → TCA Generation of Precursor Metabolites and Energy |
| MAR:NAP | Maromolecule → Nucleic Acid Processing Macromolecule Modification |
